# Phenology and function in lycopod-Mucoromycotina symbiosis

**DOI:** 10.1101/2020.09.28.316760

**Authors:** Grace A. Hoysted, Martin I. Bidartondo, Jeffrey G. Duckett, Silvia Pressel, Katie J. Field

## Abstract

*Lycopodiella inundata* is a lycophyte with a complex life cycle. The gametophytes and the juvenile, mature and retreating sporophytes form associations with Mucoromycotina fine root endophyte (MFRE) fungi, being mycoheterotrophic as gametophytes and mutualistic as mature sporophytes. However, the function of the symbiosis across juvenile and retreating sporophyte life stages remains unknown. We measured carbon-for-nutrient exchanges between *L. inundata* and MFRE across the transition from newly emerging sporophytes to mature sporophytes and in retreating adult sporophytes. We show MFRE fungi play distinct functional roles at each plant life stage, with evidence of bidirectional exchange of plant C for fungal acquired nutrients (N and P) between mature adult and retreating adult sporophytes and fungi, but no transfer of plant C to fungi and little fungal-acquired nutrient gain in juvenile sporophytes. Furthermore, we show that these functional stages correspond with different cytologies of colonisation. Our results show that MFRE have considerable plasticity in their interactions with the host plant which is related to the developmental stage of the host. This highlights the need for further research into symbiotic fungal function across plant life histories.

## Introduction

Lycopods represent a significant node on the land plant phylogenetic tree, being the earliest divergent extant tracheophyte lineage (Kenrick, 1994) and marking the transition from non-vascular to vascular plants. Several lycophytes (*Huperzia, Lycopodium, Lycopodiella* and *Phylloglossum*; Supplementary Fig. **1a**) possess an “alternation of generations” lifecycle (Kenrick, 1994) which features fully independent gametophyte (haploid) and dominant sporophyte (diploid) generations (Haufler et al, 2016; Supplementary Fig. **1b**). In nature, all members of the Lycopodiaceae require mycorrhizal symbionts for growth and for the production of gametes (Winther and Friedman, 2008). These fungal symbionts are of particular interest as they are reported to be present across both free-living generations of the plants: from the gametophyte to the young sporophyte (protocorm), while still attached to the gametophyte, through to the mature sporophyte (Bierhorst, 1971; Winther and Friedman, 2008).

Initially, it was thought that the fungal symbionts of the Lycopodiaceae were arbuscular mycorrhizal (AM)-like with unique “lycopodioid” features (Schmid and Oberwinkler, 1993). However, a recent global analysis of over 20 lycopod species determined that many form symbioses with both AM-forming Glomeromycotina fungi and Mucoromycotina “fine root endophyte” (MFRE) fungi, with *Lycopodiella inundata* being the only lycopod species found to engage in exclusive associations with MFRE partners (Rimington et al. 2015). MFRE, previously classified as the AM species *Glomus tenue*, have recently been reclassified as belonging within the Mucoromycotina (Orchard et al, 2017a, b) and renamed as *Planticonsortium tenue* (Walker et al. 2018). Emerging evidence suggests that, in contrast to the majority of studies on MFRE which have so far focussed primarily on the role of the fungal partners in phosphorus (P) transfer to host plants (Orchard et al, 2017a), and in contrast to AM fungi that are predominantly involved in P uptake (Smith & Read, 2008), MFRE partners play a significant role in plant nitrogen (N) assimilation (Hoysted et al, 2019; Field et al, 2019).

Mycorrhizal functioning in plants with alternating generations, such as *L. inundata*, is complex and poorly understood with the only published research to date focussing on instantaneous measurements on a single life history stage, e.g. photosynthetic sporophytes of *Ophioglossum* associating with AMF (Field et al, 2015; Suetsugu et al, 2020). To date, only one study has dissected the symbiotic function of MFRE in *L. inundata,* or indeed in any vascular plant (Hoysted et al, 2019); however, like other studies investigating mycorrhizal function, experiments were limited to actively growing, photosynthetic adult sporophytes with erect fertile stems and thus provide only a snapshot in time of symbiotic function in a perennial plant. Given that MFRE have been reported to be present at each life stage of *L. inundata* – from the subterranean gametophyte to the retreating adult sporophyte (Hoysted et al, 2019), these plants provide a unique opportunity to understand symbiotic function and enhance our knowledge of MFRE, not only in a vascular plant, but one with a complex lifecycle.

We used a combination of isotope tracers and cytological analyses to investigate how MFRE fungal morphology and function may change across the transition from newly emerging, juvenile sporophytes to retreating adult sporophytes of *L. inundata*, how MFRE function changes as plants become photosynthetic and how the loss of photosynthetic capacity of *L. inundata* may affect MFRE-acquired nutrient assimilation in retreating sporophytes.

## Materials and Methods

We collected *Lycopodiella inundata* (L.) gametophytes and sporophytes at three different life stages (Figure 2a-c, Figure S1b) from Thursley National Nature Reserve, Surrey, UK (SU 90081 39754) in spring and late summer, 2017. Using the methods of Hoysted et al, (2019), we quantified carbon-for-nutrient exchange between *L. inundata* and MFRE symbionts. ^33^P-labelled orthophosphate and ^15^N-labelled ammonium chloride were used to trace nutrient flow from MFRE-to-plant for each of the *L. inundata* life stages collected. We simultaneously traced the movement of carbon from plant-to-MFRE by generating a pulse of ^14^CO_2_ and quantifying the activity of extraradical MFRE hyphae in the surrounding soil using sample oxidation (307 Packard Sample Oxidiser, Isotech, Chesterfield, UK) and liquid scintillation (see Supplementary Information for details).

*L. inundata* gametophytes, juvenile sporophytes (up to 7 leaves, remnants of protocorm, rhizoids) and roots of mature and retreating adult plants (both wild and experimental), were either stained with trypan blue (Brundrett et al, 1996), fixed and embedded in Spur’s resin following Hoysted et al (2019), or processed for scanning electron microscopy (SEM) (Hoysted et al. 2019), within 48 hrs of collection (Orchard et al, 2017c).

Fungal symbionts from root samples of experimental plants were identified using molecular fungal identification methods as per Hoysted et al. (2019; see Supplementary Information for details) with MFRE being detected in each life stage (GenBank/EMBL accession numbers: MK673773-MK673803).

## Results and Discussion

Our data show MFRE fungi play distinct functional roles at each life stage of *L. inundata,* with evidence of bidirectional exchange of plant C for fungal acquired nutrients (N and P) between mature adult and retreating adult sporophytes and fungi, but no transfer of plant C to fungi and little fungal-acquired nutrient gain in juvenile sporophytes. Furthermore, we show that these functional stages correspond with different cytologies of colonisation across the *L. inundata* life cycle. Considered alongside the results of studies in other plants with complex life cycles (Roy et al., 2013; Gonneau et al., 2014; Suetsugu et al., 2018), our results emphasise the importance of investigating symbiotic fungal function across plant life histories.

### C-for-nutrient exchange between L. inundata and MFRE across life stages

*L. inundata* forms associations with MFRE fungi in each stage of its life cycle (Rimington et al, 2015; Hoysted et al, 2019) and previous research in mature sporophytes has demonstrated that these associations represent nutritional mutualisms, akin to AM fungal associations in other vascular plants (Hoysted et al, 2019). However, despite there being copious MFRE colonisation within juvenile sporophytes (Figure 2d-f), we found no transfer of plant C to MFRE (Fig. 1a, b; Table S3,4) even though green leaves were present with potential photosynthetic capabilities. In contrast, transfer of C from plants to MFRE in both the mature and retreating adult sporophyte growth stages was evident (Fig. 1a, b; Table S3,4), with ~2.4 times the amount of C being transferred from the plant to MFRE in mature adult sporophytes compared to retreating adult sporophytes, although this difference was not significant (Mann-Whitney U = 142.000, *P* = 0.144).

**Figure 1.**
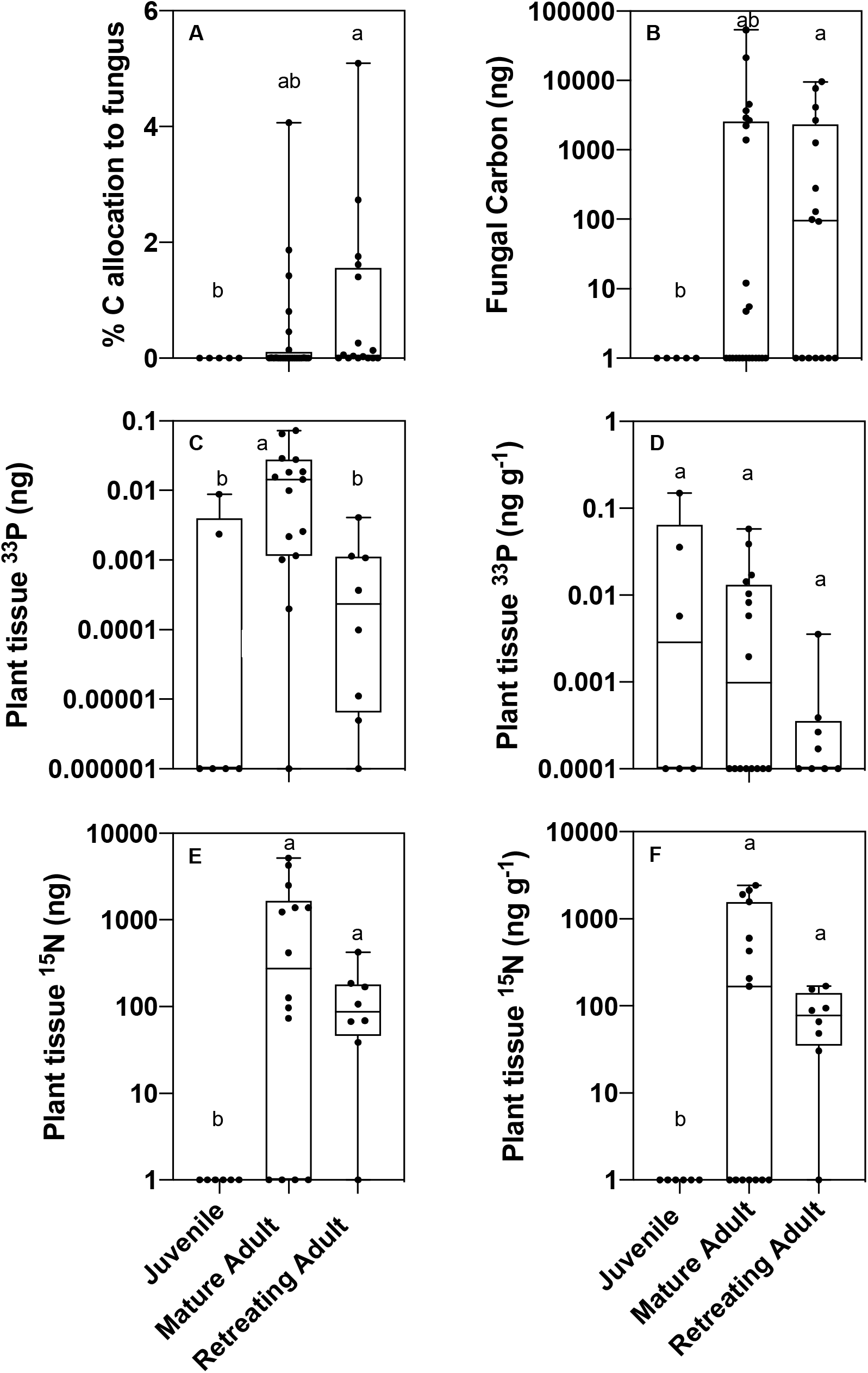
Carbon-for-nutrient exchange between *Lycopodiella inundata* sporophytes (juvenile, mature adult and retreating adult) and Mucoromycotina fine root endophyte (MFRE). (a) Percentage allocation of plant-derived carbon to fungi within soil cores; (b) total measured plant-fixed carbon transferred to MFRE in soil by lycophyte sporophytes; (c) total plant ^33^P content (ng) and (d) tissue concentration (ng g^−1^) of fungal acquired ^33^P in juvenile, mature adult and retreating adult *L. inundata* plants; (e) total tissue ^15^N content (ng) and (f) concentration (ng g^−1^) of fungal-acquired ^15^N in lycophyte sporophytes. In all panels, error bars show minimum to maximum data points. Different letters represent where *P* < 0.05 (Mann-Whitney U test). The absence of a bar denotes no transfer of carbon or nutrients. In panels (a) and (b), *n*=5, *n*=24, *n*=16; in panels (c) and (d), *n*=5, *n*=15, *n*=8; in panels (e) and (f), *n*=6, *n*=15, *n*=8 for juvenile, mature adult and retreating adult sporophytes, respectively.

**Figure 2.**
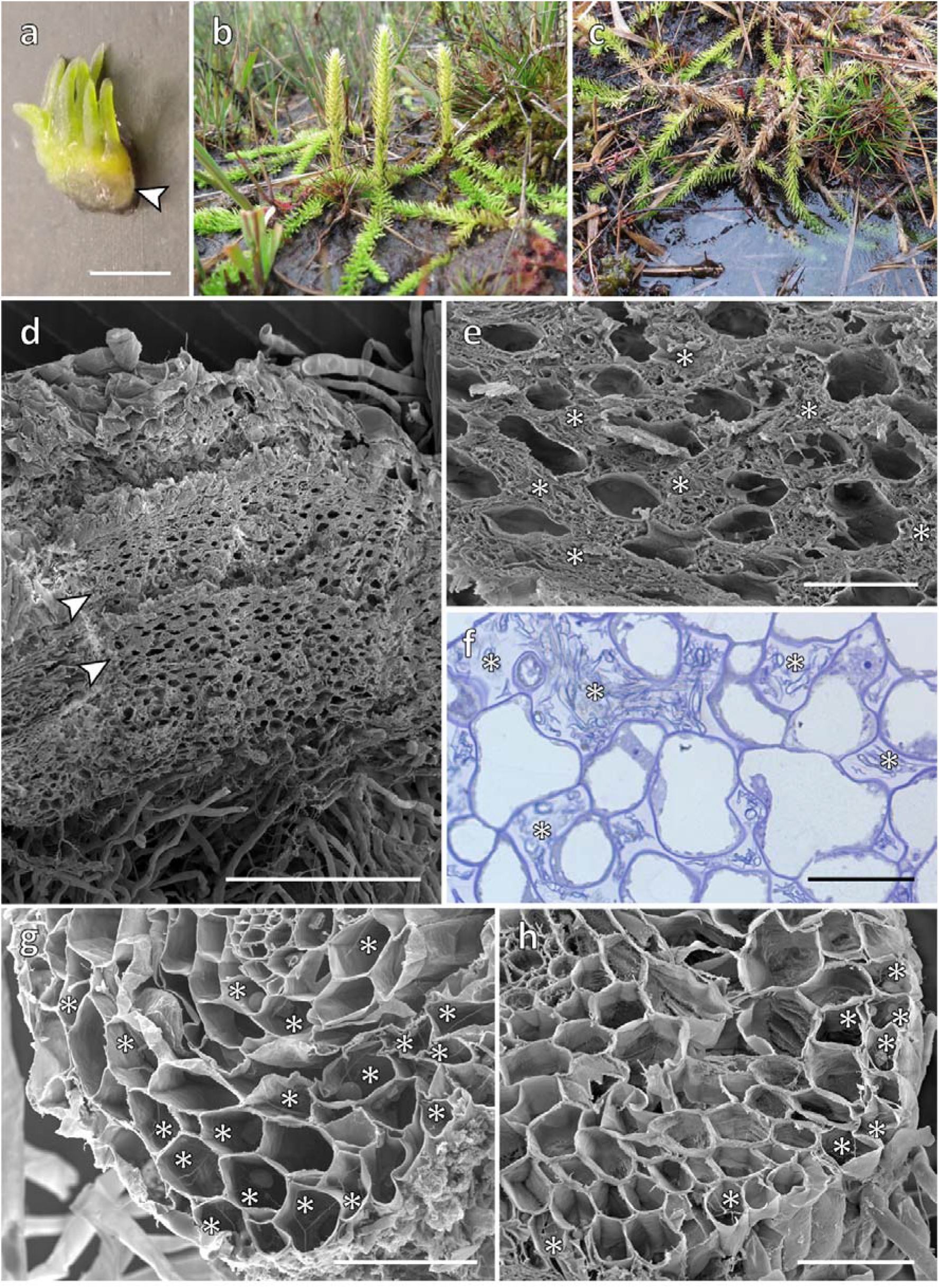
Patterns of Mucoromycotina fungal colonisation in *L. inundata*. Scanning electron micrographs, except (a-c) digital camera images and (f) light micrograph of toluidine blue stained semi-thin section. (a - c) Life stages of *L. inundata* analysed in this study. (a) Example of juvenile sporophyte at the developmental stage used in our isotope tracer experiments; the sporophyte is no longer attached to the gametophyte, has up to seven leaves and remnants of protocorms (yellowish, arrowed) with copious rhizoids emerging from the ventral side. (b, c) *L. inundata* at Thursely Common; (b) mature adult sporophytes in summer and (c) retreating adult sporophytes in spring, note the partially submerged creeping stems. (d-f) Protocorms of juvenile sporophytes are almost completely filled by an extensive system of intercellular spaces (d, arrowed), which is packed with swollen, pseudoparenchymatous, mostly collapsed hyphae (e, *, f, *). (g). Transverse section of root of mature sporophyte of *L. inundata* showing extensive fungal colonisation (*). (h) In the roots of retreating sporophytes the fungus is largely confined to the epidermal and outermost cortical layers (h,*). **Scale bars:**(a) 1 mm; (d) 500 μm; (g, h) 100 μm; (e, f) 50 μm.

Winther and Friedman (2008) suggested a form of parental nurture may occur in lycopods with achlorophyllous subterranean gametophytes, such as *L. inundata*, where fidelity of fungal partners and shared mycelial networks between generations allow autotrophic sporophytes to supply the small but critical amounts of carbohydrates required to support heterotrophic gametophytes (Leake et al, 2008). Our findings corroborate this idea of intergenerational support, with adult and retreating sporophytes transferring C to MFRE partners and C transfer by juveniles being undetectable.

Movement of ^33^P from MFRE associates was detected in all *L. inundata* plants tested, although the amounts transferred varied among growth stages (Fig. 1c, d; Table S2), with juvenile *L. inundata* sporophytes receiving approximately 10-fold less 33P from their fungal partner compared to mature adult *L. inundata* sporophytes (Mann-Whitney U= 13.000, *P* = 0.012, Fig. 1c). However, there was no significant difference in the amounts of ^33^P received from MFRE between mature adult sporophytes and juvenile sporophytes when above-ground plant tissue ^33^P content was normalised to plant biomass (Mann-Whitney U = 45.000, *P* = 0.813, Fig. 1d). In addition to ^33^P, significant amounts of ^15^N were transferred from MFRE to the shoots of mature and retreating adult *L. inundata* sporophytes (Fig. 1e, f; Table S2). Mature adult sporophytes received ~9 times more ^15^N from MFRE compared to retreating ones. However, there was no ^15^N transferred from MFRE to any of the juvenile sporophytes tested (Fig. 1e, f; Table S2).

Although there was little-to-no exchange of plant-fixed C for fungal-acquired nutrients in juvenile sporophytes, we observed abundant bi-directional exchange of carbon for ^33^P and ^15^N between the mature adult sporophyte of *L. inundata* and MFRE fungi (Fig. 1a-f; Table S2-4). These results are similar to those of a previous investigation into the function of AMF symbionts of green sporophytes of the fern *Ophioglossum vulgatum*, also defined by a characteristic alternation of generations (Field et al, 2015), which showed mutualistic exchange of plant fixed carbon for nutrients between symbionts.

### Changing patterns of colonisation

SEM results confirm distinct differences in fungal colonisation between gametophytes, juvenile sporophytes and roots of adult plants. Colonisation of the protocorm of newly developing sporophytes, which remain attached to the gametophyte (Fig. S2a-c), occurs *de novo*, with no evidence of the fungal symbiont crossing the gametophyte-sporophyte junction (placenta) (Fig. S2d). Fungal colonisation in newly developing sporophytes is both intra- and intercellular (Fig. S2e) and, like in the gametophytes, consists of thin (>2 μm in diameter), branching hyphae with small intercalary and terminal vesicles (Fig. S2d), typical of MFRE colonisation. As the young sporophytes develop the intercellular hyphae enlarge, reaching diameters well in excess of 3 μm, while the vesicles disappear (see Hoysted et al, 2019). By the time young sporophytes have reached the developmental stage used in our isotope tracer experiments (up to seven leaves, remnants of protocorm, rhizoids and no or rarely one newly developing rootlet) (Fig. 2a), the system of large, mucilage-filled intercellular spaces almost completely fills the remnants of the protocorm (Fig. 2d) and is packed with pseudoparenchymatous hyphal masses (Fig. 2e), which are mostly collapsed (Fig. 2f). Roots of actively growing (Fig. 2g) and retreating (Fig. 2h, 2Sf, g) adult plants both display the same cytology of colonisation, consisting of intracellular thin hyphae and vesicles (Fig. S2f, g) (Hoysted et al, 2019), however in the latter the fungus is largely confined to the epidermal and outermost cortical layers (Fig. 2h).

MFRE fungi have a distinct zonation in the gametophytes and protocorms of newly developed *L. inundata* sporophytes consisting of an intracellular phase of colonisation characterised by fine hyphae with small swelling/vesicles (and, in the gametophyte only, also hyphal coils with larger vesicles – see Hoysted et al, 2019) and an intercellular phase where the fungus proliferates in the system of mucilage-filled intercellular spaces forming masses of large pseudoparenchymatous hyphae that eventually collapse and degenerate (Hoysted et al, 2019). This colonisation is the same as that reported in other lycopod gametophytes and protocorms (Schmid and Oberwinkler, 1993; Duckett and Ligrone, 1992) and strikingly similar to that described in the earliest diverging Haplomitriopsida liverworts *Treubia* and *Haplomitrium* (Duckett et al, 2006; Carafa et al, 2003), the only two liverwort genera known to date to be colonised exclusively by MFRE fungi (Bidartondo et al, 2011; Field et al, 2015; Rimington et al, 2020). In *Treubia* and *Haplomitrium*, the intracellular fungal swellings or ‘lumps’ are relatively short-lived; it has been suggested that these structures are involved in active metabolic interactions with the host cells (Carafa et al, 2003) and that their eventual collapse and lysis may also provide nutrients, such as nitrogen, to the host plant (Duckett et al, 2006).

The MFRE fungal colonisation in the roots of adult sporophytes is only intracellular and consists of fine aseptate hyphae with intercalary and terminal swellings/vesicles but without arbuscules (Hoysted et al, 2019). It is possible that the small swellings/vesicles may play an important role in host-fungus physiological relationships, as it has been suggested for Haplomitriopsida liverworts (Duckett et al, 2006; Carafa et al, 2003). Further studies are urgently needed to determine the functional role of the diverse structures produced by MFRE in the different stages of *Lycopodiella's* life cycle, and indeed other plants. In retreating sporophytes, fungal colonisation appears much reduced compared to fully photosynthesising sporophytes, being mostly restricted to the outermost cortical layers (Fig. 2h). This may explain why retreating sporophytes receive smaller amounts of N and P from their fungal symbionts (Fig. 1c-f).

### Intergenerational support by MFRE in L. inundata

Previous descriptions of *Lycopodiella* have highlighted the crucial role played by symbiotic fungi in the continued growth of the gametophyte; growing green portions of older gametophytes of *L. alopecuroides* were often observed to be embedded in older, yellow portions with abundant fungal hyphae (Koster, 1941). Coupled with the absence of C and N transfer between *L. inundata* and MFRE fungi in the juvenile sporophyte in our experiments (Fig. 1a.,b,e,f; Table S2-S4) this may suggest the presence of intergenerational support between alternating life stages whereby later life stages need to be present to transfer essential nutrients and nurture younger plants.

In our experiments, the juvenile sporophytes were sustained throughout the experimental period despite the apparent lack of photosynthetic carbon being transferred from plant-to-fungus and without hyphal connections to mature sporophytes. It is possible that residual carbon reserves within the sporophyte tissues were mobilised and used for plant growth and allocation to fungi and recent photosynthates restricted for use only in plant tissues, suggestive of there being intricate temporal dynamics in allocation of carbon resources to fungal partners in this key transitional stage. Alternatively, the presence of collapsed and degenerating pseudoparenchymatous hyphal masses filling the extensive system of intercellular spaces in the remnants of protocorms may suggest a different scenario. This juvenile sporophytic stage just precedes root development and therefore formation of a mycorrhizal association *sensu stricto* between *Lycopodiella* sporophytes and MFRE symbionts. It is likely that very early stages of sporophyte development are, like the gametophytes, completely, or largely mycoheterotrophic, as fungal colonisation is ubiquitous and extensive in their subterranean protocorms with only the apical parts of newly developing, green leaves emerging above the ground. It is therefore possible that juvenile sporophytes just prior to root development maintain a partially mycoheterotrophic lifestyle, the masses of collapsed and degenerating intercellular hyphae releasing nutrients that support early sporophyte development. Further investigations are now required that include structural and functional assessment of subterranean gametophytes associating with MFRE fungi.

## Conclusion

This investigation represents the first functional assessment of fungal symbiosis across the changing phenology of the marsh clubmoss, *L. inundata*. We show that MFRE fungi play critical and distinct functional roles across different developmental stages and that these correspond with different cytologies of colonisation. Our results show that MFRE have considerable plasticity in their interactions with plants which appears to relate to the developmental stage of the host and is suggestive of intergenerational support between sporophytes and gametophytes via shared MFRE symbionts.

## Acknowledgements

We gratefully acknowledge support from the NERC to KJF, SP, SS (NE/N00941X/1; NE/S009663/1) and MIB (NE/N009665/1). KJF is funded by a BBSRC Translational Fellowship (BB/M026825/1) and a Philip Leverhulme Prize (PLP-2017-079). We thank James Giles (Natural England) for field support, Julia Masson and the RSPB for access to the Norfolk site, and The Species Recovery Trust for access to the Dorset site.

## Author Contributions

KJF, SP, MIB and JGD conceived and designed the investigation. SP and JGD collected plant material. GAH undertook physiological analysis. SP undertook the cytological analysis. GAH led the writing; all authors discussed results and comments on the article. GAH agrees to serve as the author responsible for contact and ensure communication.

