## Supplementary File for "Phenology and function in lycopod-Mucoromycotina symbiosis"

Supplementary Figure 1a

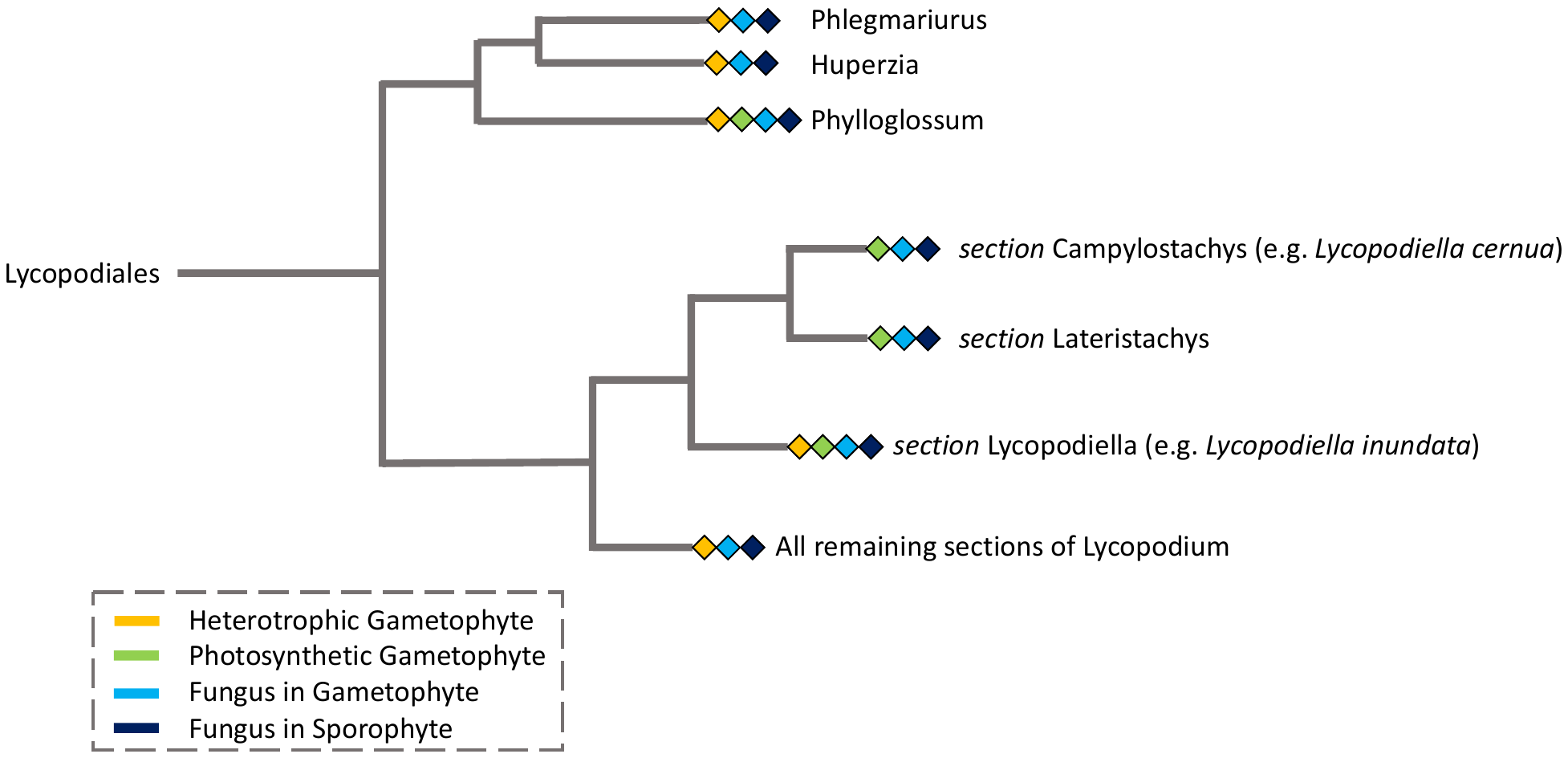

Supplementary Figure 1b

Mature Adult Sporophyte

Retreating Adult

Sporophyte

Spore

Mucoromycotina FRE

Gametophyte

Juvenile Sporophyte

**Autotrophy**

**Mycoheterotrophy**

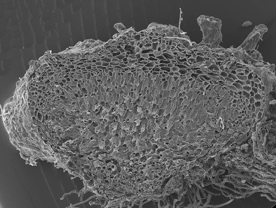

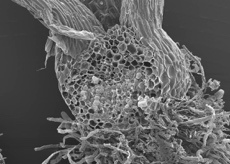

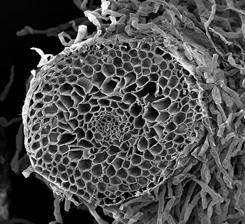

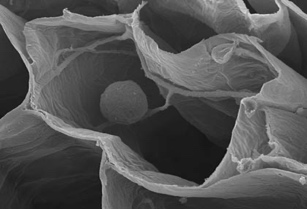

Mucoromycotina FRE

**Supplmentary figure 1. Land plant phylogeny of the Lycopodiales and life cycle of *Lycopodiella inundata*.** (a) Land plant phylogeny showing key nodes within the Lycopodiales alongside gametophyte and sporophyte stages within each genus (Field et al., 2016). (b) Schematic diagram showing key stages of the life cycle in *Lycopodiella inundata*. Spore germination gives rise to a short-lived subterranean gametophyte, which is nutritionally completely reliant on a fungal symbiont (mycoheterotrophy). Following fertilisation, the newly developing sporophyte, which lacks roots, retains a partially mycoheterotrophic lifestyle until it develops into an autotrophic adult plant, able to generate its own carbohydrates. During summer, the adult sporophyte consists of vegetative creeping stems tightly appressed to the soil and erect fertile stems. During winter and spring the vegetative stems are often under water or ice.

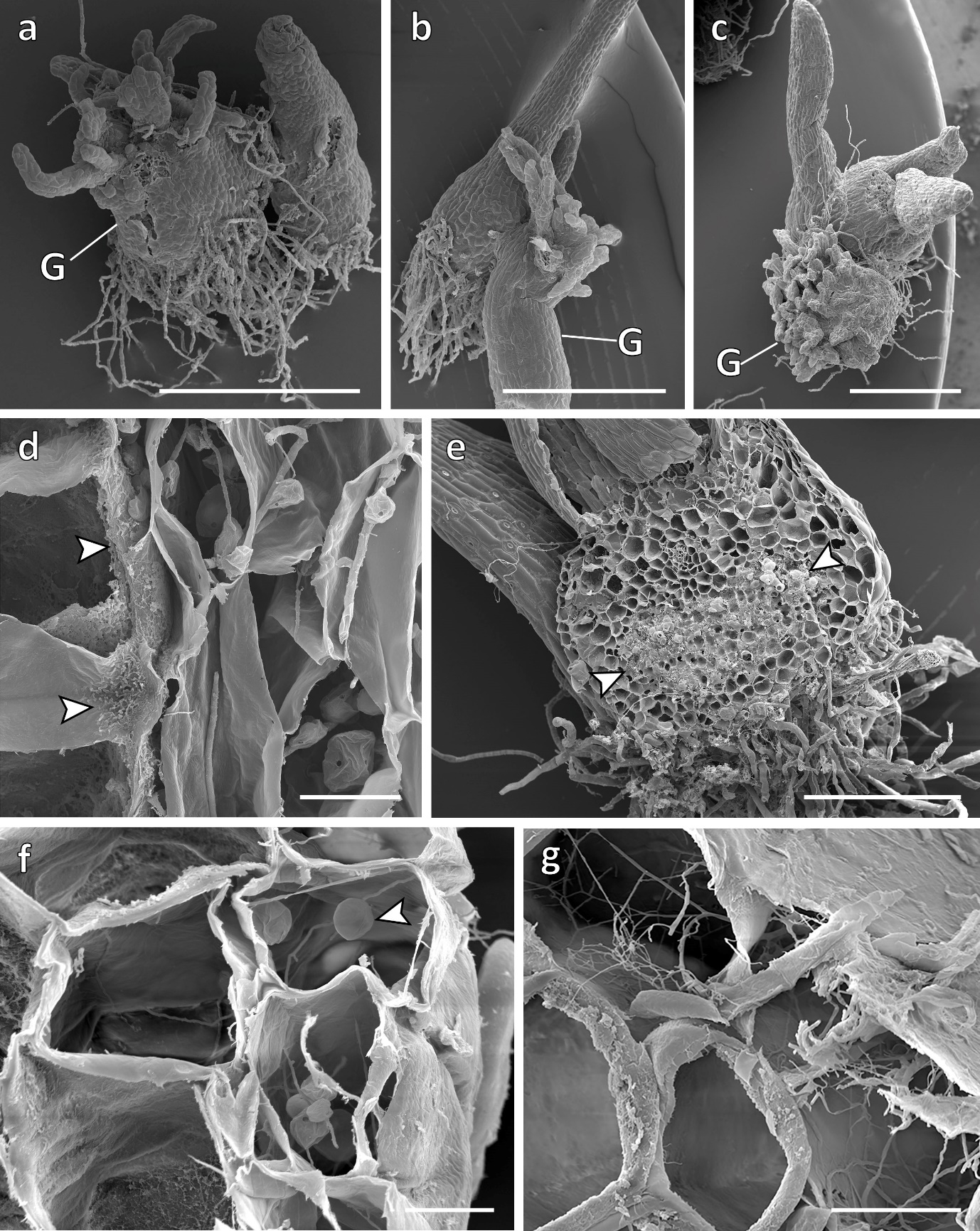

**Supplementary figure 2. Patterns of fungal colonisation in *L. inundata*.** (a-c) Newly developing sporophytes of *L. inundata* still attached to the gametophytes (G). (d) The gametophyte fungus does not cross the gametophyte-sporophyte junction (arrowed), identifiable by its numerous wall ingrowth. (e) Newly developing sporophyte showing zone of intercellular fungal colonisation (arrowed). (f, g) In the adult sporophyte roots, shown here in retreating adults, colonisation is exclusively intracellular and, as in the gametophytes and young sporophyte protocorms, consists of branching fine hyphae with intercalary and terminal small vesicles (arrowed in f, also note the collapsed vesicles in the cell below) **Scale bars:** (a-c) 1 mm; (e) 500 µm; (d, f, g) 20 µm.

**Supplementary Table 1:** Summary of differences in Mucoromycotina fine root endophyte functionality (Z value from Mann Whitney U and Kruskal Wallis H value) between juvenile, mature adult and retreating adult sporophytes of *Lycopodiella inundata*. Bold numbers represent significant values.

|  |  |  | *P-*value | | |
| --- | --- | --- | --- | --- | --- |
|  | Mann-Whitney U | Z value | Juvenile v. mature adult | Juvenile v. retreating adult | Mature adult v. retreating adult |
| Percentage of carbon allocation (%) | 32.500 | -1.820 | 0.069 |  |  |
|  | 15.000 | -2.230 |  | **0.026** |  |
|  | 142.000 | -1.461 |  |  | 0.144 |
| Kruskal-Wallis H | 7.096 |  |  |  |  |
| Asymp. Sig | **0.029** |  |  |  |  |
| Fungal carbon in core (ng) | 32.500 | -1.820 | 0.069 |  |  |
|  | 17.500 | -2.059 |  | **0.040** |  |
|  | 174.500 | -0.516 |  |  | 0.606 |
| Kruskal-Wallis H | 4.969 |  |  |  |  |
| Asymp. Sig | 0.083 |  |  |  |  |
| Total ^33^P uptake (ng) | 13.000 | -2.058 | **0.012** |  |  |
|  | 17.000 | -0.924 |  | 0.355 |  |
|  | 17.500 | -2.744 |  |  | **0.006** |
| Kruskal-Wallis H | 10.867 |  |  |  |  |
| Asymp. Sig | **0.004** |  |  |  |  |
| [^33^P] in plant tissue (ng g^-1^) | 45.000 | -0.236 | 0.813 |  |  |
|  | 18.000 | -0.827 |  | 0.408 |  |
|  | 49.000 | -0.982 |  |  | 0.326 |
| Kruskal-Wallis H | 1.144 |  |  |  |  |
| Asymp. Sig | 0.564 |  |  |  |  |
| Total ^15^N uptake (ng) | 10.000 | -2.477 | **0.014** |  |  |
|  | 2.500 | -2.695 |  | **0.007** |  |
|  | 42.000 | -0.961 |  |  | 0.337 |
| Kruskal-Wallis H | 9.088 |  |  |  |  |
| Asymp. Sig | **0.011** |  |  |  |  |
| [^15^N] in plant tissue (ng g^-1^) | 17.500 | -1.970 | **0.049** |  |  |
|  | 2.500 | -2.695 |  | **0.007** |  |
|  | 53.500 | -0.429 |  |  | 0.688 |
| Kruskal-Wallis H | 6.614 |  |  |  |  |
| Asymp. Sig | **0.037** |  |  |  |  |

**Supplementary Table 2.** Summary of the amounts of ^15^N and ^33^P detected in static and rotated core of microcosms used during carbon-for-nutrient experiments between *Lycopodiella inundata* juvenile, mature adults and retreating adults and Mucoromycotina FRE fungi. SEM = standard error of the mean.

|  |  | **Isotope introduced to static core** | | | | **Isotope introduced to rotated core** | | | |
| --- | --- | --- | --- | --- | --- | --- | --- | --- | --- |
|  |  | **ng** | **SEM** | **ng/g^-1^** | **SEM** | **ng** | **SEM** | **ng/g^-1^** | **SEM** |
| **Mean^15^N in plant tissue** | **Juvenile** | 0 | NA | 0 | NA | 0 | NA | 0 | NA |
|  | **Mature Adult** | 1512.951112 | 428.8337579 | 1068.898435 | 273.3093604 | 310.279627 | 123.056841 | 521.568421 | 133.243177 |
|  | **Retreating Adult** | 140.1084306 | 48.60599254 | 88.68543443 | 21.99000531 | 7.360712123 | 5.037044815 | 7.01358979 | 4.82637283 |
| **Mean ^33^P in plant tissue** | **Juvenile** | 0.00224315 | 0.00156839 | 0.039163362 | 0.025311421 | 0.001199979 | 0.000303609 | 0.018268206 | 0.005124894 |
|  | **Mature Adult** | 0.032491994 | 0.011026128 | 0.022264824 | 0.004830504 | 0.00443139 | 0.001006564 | 0.01659068 | 0.0044485 |
|  | **Retreating Adult** | 0.001303603 | 0.000476245 | 0.000993662 | 0.000421648 | 0.000471753 | 9.42274E-05 | 0.000515893 | 0.000107934 |

**Supplementary Table 3**: Summary of the % amount of carbon detected in static and rotated core of microcosms used during carbon-for-nutrient experiments between *Lycopodiella inundata* juvenile, mature adults and retreating adults and Mucoromycotina FRE fungi. The average value (in bold) for the static minus rotated core column is represents the value used to produce Figure 1a in the main text

| **Carbon Allocated to Fungus** | | | | | |
| --- | --- | --- | --- | --- | --- |
|  |  | **Static Core** |  | **Rotated Core** | **Static Minus Rotated Core** |
|  | **Plant Replicate** | **%** | **Plant Replicate** | **%** | **%** |
|  | 1 | 5.549921505 | 2 | 6.52525124 | 0 |
|  | 2 | 2.093482848 | 3 | 18.86149918 | 0 |
|  | 3 | 1.702356322 | 4 | 2.850043869 | 0 |
|  | 4 | 0.433085719 | 5 | 0.780794239 | 0 |
|  | 5 | 0.317956442 | 6 | 0.68111204 | 0 |
| **Average** |  | 1.970407773 | **Average** | 5.21219623 | **0** |
| **Mature Adult**  **Sporophyte** | 1 | 0.028734233 | 1 | 0.021393887 | 0.007340347 |
|  | 2 | 0.032034103 | 2 | 0.02934499 | 0.002689113 |
|  | 3 | 0.022253088 | 3 | 0.048716375 | 0 |
|  | 4 | 0.009479151 | 4 | 0.013452419 | 0 |
|  | 5 | 0.01262442 | 5 | 0.003241787 | 0.009382633 |
|  | 6 | 0.015150077 | 6 | 0.00854962 | 0.006600456 |
|  | 7 | 0.021319134 | 7 | 0.17124069 | 0 |
|  | 8 | 0.230976317 | 8 | 6.541434471 | 0 |
|  | 9 | 1.48129678 | 9 | 0.058614498 | 1.422682283 |
|  | 10 | 9.047650886 | 10 | 7.177982219 | 1.869668667 |
|  | 11 | 2.318153038 | 11 | 9.887981329 | 0 |
|  | 12 | 0.221764154 | 12 | 0.080698028 | 0.141066126 |
|  | 13 | 0.018352876 | 13 | 0.017352255 | 0.001000622 |
|  | 14 | 0.01563522 | 14 | 0.017012946 | 0 |
|  | 15 | 0.020713834 | 15 | 0.176972838 | 0 |
|  | 16 | 0.035800798 | 16 | 0.053049036 | 0 |
|  | 17 | 0.016416467 | 17 | 0.062033129 | 0 |
|  | 18 | 0.740868564 | 18 | 1.657575604 | 0 |
|  | 19 | 4.261856746 | 19 | 0.194228682 | 4.067628064 |
|  | 20 | 0.338974679 | 20 | 1.290564524 | 0 |
|  | 21 | 0.27559024 | 21 | 0.554999977 | 0 |
|  | 22 | 0.580869344 | 22 | 0.122975253 | 0.457894092 |
|  | 23 | 0.025155723 | 23 | 0.820308424 | 0 |
|  | 24 | 1.071227043 | 24 | 0.262211095 | 0.809015948 |
| **Average** |  | 0.868454038 | **Average** | 1.21966392 | **0.366457015** |
| **Retreating Sporophyte** | 1 | 0.052165841 | 1 | 0.089781891 | 0 |
|  | 2 | 0.246865135 | 2 | 0.188011761 | 0.058853374 |
|  | 3 | 0.131509311 | 3 | 0.202741732 | 0 |
|  | 4 | 1.827429133 | 4 | 0.070924321 | 1.756504812 |
|  | 5 | 0.218964217 | 5 | 0.087276452 | 0.131687765 |
|  | 6 | 0.106030651 | 6 | 0.077765399 | 0.028265252 |
|  | 7 | 0.087433796 | 7 | 0.107967692 | 0 |
|  | 8 | 0.284634296 | 8 | 0.022820308 | 0.261813988 |
|  | 9 | 5.516404975 | 9 | 0.424638246 | 5.091766729 |
|  | 10 | 0.32291578 | 10 | 0.284457017 | 0.038458763 |
|  | 11 | 0.046422816 | 11 | 2.536998575 | 0 |
|  | 12 | 0.094223144 | 12 | 6.797492519 | 0 |
|  | 13 | 0.533424254 | 13 | 3.809266719 | 0 |
|  | 14 | 1.903249828 | 14 | 0.285611066 | 1.617638762 |
|  | 15 | 2.94128412 | 15 | 0.207659922 | 2.733624198 |
|  | 16 | 1.684305138 | 16 | 0.283626401 | 1.400678736 |
| **Average** |  | 0.999828902 | **Average** | 0.967315001 | **0.819955774** |

**Supplementary Table 4**: Summary of the total amount of carbon detected in static and rotated cores in microcosms used during carbon-for-nutrient experiments between *Lycopodiella inundata* juvenile, mature adults and retreating adults and Mucoromycotina FRE fungi. The average value (in bold) for the static minus rotated core column is represents the value used to produce Figure 1b in the main text.

| **Carbon Allocated to Fungus** | | | | | |
| --- | --- | --- | --- | --- | --- |
|  |  | **Static Core** |  | **Rotated Core** | **Static Minus Rotated Core** |
|  | **Plant Replicate** | **ng** | **Plant Replicate** | **ng** | **ng** |
| **Juvenile Sporophyte** | 1 | 2043.81579 | 1 | 2402.991016 | 0 |
|  | 2 | 2176.527077 | 3 | 19609.69669 | 0 |
|  | 3 | 2369.735289 | 4 | 3967.353628 | 0 |
|  | 4 | 1451.121022 | 5 | 2616.172469 | 0 |
|  | 5 | 1178.90384 | 6 | 2525.394974 | 0 |
| **Average** |  | 1995.753807 | **Average** | 5605.790078 | **0** |
| **Mature Adult**  **Sporophyte** | 1 | 47.17233028 | 1 | 35.12185178 | 12.0504785 |
|  | 2 | 65.7876335 | 2 | 60.26506972 | 5.522563771 |
|  | 3 | 53.59773982 | 3 | 117.3359678 | 0 |
|  | 4 | 41.42763416 | 4 | 58.79238765 | 0 |
|  | 5 | 3584.867206 | 5 | 920.5473911 | 2664.319815 |
|  | 6 | 3206.191204 | 6 | 1809.345113 | 1396.846091 |
|  | 7 | 419.0831031 | 7 | 3366.181803 | 0 |
|  | 8 | 3073.139172 | 8 | 87033.76502 | 0 |
|  | 9 | 22145.00496 | 9 | 876.2716269 | 21268.73334 |
|  | 10 | 13965.89893 | 10 | 11079.88973 | 2886.009193 |
|  | 11 | 8144.708331 | 11 | 34740.90044 | 0 |
|  | 12 | 3518.081953 | 12 | 1280.199119 | 2237.882833 |
|  | 13 | 87.19546046 | 13 | 82.44145614 | 4.754004324 |
|  | 14 | 95.20376918 | 14 | 103.5928246 | 0 |
|  | 15 | 106.8101973 | 15 | 912.554576 | 0 |
|  | 16 | 51.65069553 | 16 | 76.53515413 | 0 |
|  | 17 | 2306.275827 | 17 | 8714.756306 | 0 |
|  | 18 | 2137.705754 | 18 | 4782.776702 | 0 |
|  | 19 | 55697.53383 | 19 | 2538.344018 | 53159.18981 |
|  | 20 | 1261.377355 | 20 | 4802.390763 | 0 |
|  | 21 | 1617.382693 | 21 | 3257.181228 | 0 |
|  | 22 | 4661.925729 | 22 | 986.9715111 | 3674.954218 |
|  | 23 | 1780.229563 | 23 | 58051.8909 | 0 |
|  | 24 | 5997.396089 | 24 | 1468.021001 | 4529.375088 |
| **Average** |  | 5586.068632 | **Average** | 9464.836332 | 3826.65156 |
| **Retreating Sporophyte** | 1 | 147.1097444 | 1 | 253.1885003 | 0 |
|  | 2 | 538.3320285 | 2 | 409.9920899 | 128.3399386 |
|  | 3 | 413.6854009 | 3 | 637.7593645 | 0 |
|  | 4 | 36.12457277 | 4 | 238.253599 | 0 |
|  | 5 | 467.6907794 | 5 | 186.4158101 | 281.2749692 |
|  | 6 | 372.0327257 | 6 | 272.8576403 | 99.17508543 |
|  | 7 | 255.3499767 | 7 | 315.3191208 | 0 |
|  | 8 | 1379.914686 | 8 | 110.6334636 | 1269.281223 |
|  | 9 | 10385.15397 | 9 | 799.4216492 | 9585.732321 |
|  | 10 | 781.2225144 | 10 | 688.1801377 | 93.04237669 |
|  | 11 | 220.1257306 | 11 | 12029.83162 | 0 |
|  | 12 | 360.904756 | 12 | 26036.56874 | 0 |
|  | 13 | 492.7603828 | 13 | 3518.8796 | 0 |
|  | 14 | 3156.123412 | 14 | 473.6234622 | 2682.49995 |
|  | 15 | 8249.197292 | 15 | 582.4080908 | 7666.789202 |
|  | 16 | 4966.621576 | 16 | 836.3478635 | 4130.273713 |
| **Average** |  | 373.5339495 | **Average** | 836.4650446 | **260.603185** |

**Materials and Methods**

*Plant material*

*Lycopodiella inundata* (L.) gametophytes and sporophytes at three different life stages (juvenile: up to seven leaves, with remnants of protocorm, rhizoids, and none or rarely with one newly developing rootlet), mature adult (creeping vegetative plants with erect fertile stems) and retreating adult (creeping vegetative plants that withstand periodic submersion, Figure 2a-c) were collected from Thursley National Nature Reserve, Surrey, UK (SU 90081 39754) in spring (retreating adults) and late summer (young and mature sporophytes) 2017. As *L. inundata's* minute (only 1-2 mm in length), subterranean gametophytes and newly developing sporophytes are non-tractable for isotope tracing experiments, we used juvenile sporophytes to represent an intermediate stage between alternating generations (Figure 2a). The *L. inundata* sporophytes of all stages were planted directly into pots (90 mm diameter x 85 mm depth) containing acid-washed silica sand. Soil surrounding plant roots was left intact and pots were weeded regularly to remove other plant species. Plants were kept well-watered.

*Growth conditions*

Based on the methods of Field et al. (2012), three plastic cores with a 10 μm nylon mesh-covered cylindrical window, were inserted into the substrate within each experimental pot. Two of the cores were filled with a homogenous mixture of acid-washed silica sand, compost (Petersfield No.2, Leicester, UK) and native soil gathered from around the roots of wild plants (in equal parts making up 99% of the core volume) and fine-ground tertiary basalt (1% core volume) to act as fungal bait (Field et al, 2012). The third core was filled with glass wool to allow below-ground gas sampling throughout the ^14^C-labelling period to monitor below ground respiration.

The experimental microcosms were maintained in a controlled environment chamber (Micro Clima 1200, Snijders Labs, The Netherlands) with settings chosen to simulate the plant’s natural environment in summer: 70% RH, 16 hr/8 hr day/night, 20°C day/ 15°C night, 100 µmol m^-2^ s^-1^ irradiance and 440 ppm atmospheric [CO_2_]. Plants were acclimated to chamber/growth regimes for four weeks for mature adult and retreating adult sporophytes (Hoysted et al, 2019) or ten weeks in the case of juvenile sporophytes to allow establishment of mycelial networks within cores and confirmed by hyphal extraction from soil and staining with trypan blue (Brundrett et al, 1996). Additionally, roots were stained to detect the presence of fungi based on the methods of Brundrett et al (1996).

*Molecular identification of fungal symbionts*

Molecular identification of fungal symbionts as Mucoromycotina in all *L. inundata* sporophytes was carried out as described previously in Hoysted et al (2019). All plants were processed for molecular analyses within one week of collection. Genomic DNA extraction and purification from all specimens and subsequent amplification, cloning and sequencing were performed according to methods from Rimington et al. (2015). The fungal 18S ribosomal rRNA gene was targeted using the fungal primer set NS1/EF3 and a semi-nested approach with Mucoromycotina- and Glomeromycotina-specific primers described in Desirò et al. (2014), was used for experimental *L. inundata* plants. Resulting partial 18S rDNA sequences were edited and preliminarily identified with BLAST in Geneious v. 8.1.7 (Kearse et al, 2012). Chimeric sequences were detected using the UCHIME2 algorithm (Edgar, 2016) in conjunction with the most recent non-redundant SSU SILVA database (SSU Ref NR 132, December 2017, www.arb-silva.de). Sequences identified as Mucoromycotina sp. were aligned with MAFFT prior to removing unreliable columns using the default settings in GUIDANCE2 (http://guidance.tau.ac.il). The best-fit nucleotide model for phylogenetic analysis was calculated using Smart Model Selection (Kumar et al. 2016). Maximum Likelihood (ML) with 1,000 replicates was performed using PhyML 3.0 (Guindon and Gascuel, 2003). Bayesian inference analysis was conducted in Mr Bayes version 3.2.6 (Ronquist and Huelsenbeck, 2003) with four Markov chain Monte Carlo (MCMC) strands and 10^6^ generations.

*Cytological analyses*

*Lycopodiella inundata* gametophytes, young sporophytes and roots of mature and retreating adult plants (both wild and experimental), were either stained with trypan blue (Brundrett et al, 1996), fixed and embedded in Spur’s resin following Hoysted et al. (2019) or processed for scanning electron microscopy (SEM) within 48 hrs of collection (Orchard et al, 2017c). For trypan blue staining, plant tissues were cleared in 5% KOH at 90^o^C for 3 h, acidified in 2% HCl for 1-2 min, stained in 0.05% trypan blue, de-stained in 50% glycerin for 24 h before viewing under a light microscope. For SEM, tissues were fixed in 3% glutaraldehyde, dehydrated through an ethanol series, critical-point dried using CO_2_ as transfusion fluid, sputter coated with 20 nm palladium-gold and viewed using a FEI Quanta scanning electron microscope (FEI, Hillsboro, OR, USA). Specimens embedded in resin were sectioned (0.5 µm thick sections) with a diamond histo-knife, stained with 0.5% toluidine blue and photographed under a Zeiss Axioscope light microscope equipped with a MRc digital camera.

*Quantification of C, ^33^P and ^15^N fluxes between lycophytes and fungi*

Following the four-week acclimation period for mature adult and retreating adult sporophytes, 100 μl of an aqueous mixture of ^33^P-labelled orthophosphate (specific activity 111 TBq mmol^-1^, 0.3 ng ^33^P added; Hartmann analytics, Braunschweig, Germany) and ^15^N-ammonium chloride (1mg ml^-1^, 0.1 mg ^15^N added; Sigma, Dorset, UK) was introduced into one of the soil-filled mesh cores in each pot through the installed capillary tube (Field et al 2016; Hoysted et al 2019). For juvenile sporophytes, 100 μl of ^33^P-labelled orthophosphate (specific activity 111 TBq mmol^-1^, 0.15 ng ^33^P added) and ^15^N-ammonium chloride (0.5 mg ml^-1^, 0.05 mg ^15^N added) was added. In half of the pots, cores containing isotope tracers were left static to preserve direct hyphal connections with the sporophytes. Fungal access to isotope tracers was limited in the remaining half of the pots by rotating isotope tracer-containing cores through 90°, thereby severing the hyphal connections between the plants and core soil. These were rotated every second day thereafter, thus providing a control treatment that allows us to determine non-mycorrhizal microbial contribution. Assimilation and transfer of ^33^P tracer from fungi into above-ground plant material was monitored using a hand-held Geiger counter held over the plant shoots daily.

At detection of peak activity of ^33^P in above-ground plant tissues (21 days after the addition of the ^33^P and ^15^N tracers), the tops of the labelled cores were sealed with plastic caps and anhydrous lanolin and the glass wool cores were sealed with rubber septa (SubaSeal, Sigma, Dorset, UK). Each pot in the three separate experiments was sealed into a 3.5 L, gas-tight labelling chamber and 2 ml 10% lactic acid was added to 30 μl NaH^14^CO_3_ (specific activity 1.998 GBq/mmol^-1^ for mature adult sporophytes, 1.621 GBq/mmol^-1^ for retreating sporophytes and 1.85 GBq/mmol^-1^ for juvenile sporophytes; Hartmann Analytics, Braunschweig, Germany) in a cuvette within the chamber prior to illumination at 0800 (Hoysted et al, 2019), releasing a 1.1 MBq pulse of ^14^CO_2_ gas. Pots were maintained under growth chamber conditions (see above), and 1 ml of gas was sampled after 1 hour and every 1.5 hours thereafter. Below-ground respiration was monitored via gas sampling from within the glass-wool filled core after 1 hour and every 1.5 hours thereafter for ~16 hours.

*Plant harvest and sample analyses*

Upon detection of peak below-ground ^14^C flux (~16 hrs), plants and soils were separated, freeze-dried, weighed and homogenised. The ^33^P activity in plant and soil sub-samples was quantified by acid digestion using 10-50 mg of homogenised plant and soil materials (in the case of juvenile sporophytes, 0.5-5 mg shoots were used). Samples were digested in 500 μl of concentrated H_2_SO_4_ and heated to 365°C for 15 min. Then, 50 μl H_2_O_2_ was added to each sample when cool. Samples were reheated to 365°C for one minute, producing a clear digest solution which was then cooled and diluted to 5 ml with distilled water. One ml of each diluted digest was then added to 10 ml of the scintillation cocktail Emulsify-safe (Perkin Elmer, Beaconsfield, UK) before liquid scintillation counting (Tricarb 2100TR liquid scintillation analyser, Isotech, Chesterfield, UK). The quantity of ^33^P tracer that was transferred to the plant by its fungal partner was then calculated using previously published equations (Cameron et al, 2007). Total ^33^P in plants without access to the tracer through core rotation (i.e. assimilated through alternative soil non-mycorrhizal microbial P-cycling processes and/or diffusion from core) was subtracted from the total ^33^P in plants with access to the core contents via intact fungal hyphal connections to give fungal acquired ^33^P.

Between two and four mg of freeze-dried, homogenised plant tissue was weighed into 6 x 4 mm^2^ tin capsules (Sercon Ltd. Irvine, UK) and ^15^N abundance was determined using a continuous flow IRMS (PDZ 2020 IRMS, Sercon Ltd. Irvine, UK). Air was used as the reference standard, and the IRMS detector was regularly calibrated to commercially available reference gases. The ^15^N transferred from fungus to plant was then calculated using equations published previously (Field et al, 2016). Total ^15^N in plants without access to the isotope because of broken hyphal connections between plant and core contents was subtracted from total ^15^N in plants with intact hyphal connections to the mesh-covered core to estimate fungal-acquired ^15^N.

The ^14^C activities of plant and soil samples were quantified through sample oxidation (307 Packard Sample Oxidiser, Isotech, Chesterfield, UK) followed by liquid scintillation. Total C (^12^C + ^14^C) fixed by the plant and transferred to the fungal network was calculated as a function of the total volume and CO_2_ content of the labelling chamber and the proportion of the supplied ^14^CO_2_ label fixed by plants. The difference in total C between the values obtained for static and rotated core contents is considered equivalent to the total C transferred from plant to symbiotic fungus within the soil core, noting that a small proportion will be lost through soil microbial respiration. The total C budget for each experimental pot was calculated using equations from Cameron et al*.* (2006).

*Statistics*

Interactions between plants and Mucoromycotina FRE fungi and the C, ^33^P and ^15^N fluxes were tested using non-parametric statistical tests. Data were originally checked for homogeneity and normality. Where assumptions for parametric tests were not met, data were transformed using log_10_. Assumptions for parametric tests following transformation were still not met, therefore a Mann Whitney U and Kruskal Wallis statistical test was performed. All statistics were performed using SPSS v24 (IBM, Armonk, New York, USA). Mean values are given with standard error mean (SEM) for error bars (Supplementary Information Table 1).
